# A longitudinal study on morphogenetic diversity of pathogenic *Rhizoctonia solani* from sugar beet and dry beans of western Nebraska

**DOI:** 10.1101/635482

**Authors:** Saurav Das, Tammy Plyler-Harveson, Dipak K. Santra, Robert M. Harveson, Kathy A. Nielsen

**Author notes:** corresponding author: Dr. Dipak Santra, Associate Professor, Panhandle Research and Extension Centre, University of Nebraska-Lincoln, USA.

## Abstract

Root and stem rot caused by *Rhizoctonia solani* is a serious fungal disease of sugar beet and dry bean production in Nebraska. Objective was to characterize morpho-genetic diversity of 38 *Rhizoctonia solani* isolated from sugar beet and dry beans fields in western Nebraska over 10 years. Classical morphological features and ISSR marker was used to study the morphogenetic diversity. Fungal colonies were morphologically diverse in shapes, aerial hyphae formation and colony, sclerotia color. Marker analysis using nineteen polymorphic ISSR marker showed polymorphic bands ranged from 15 - 28 with molecular weight 100bp to 3kb. Polymorphic loci ranged from 43.26 – 92.88%. *Nei* genetic distance within the population was ranged 0.03 –0.09 and Shannon diversity index varied from 0.24 – 0.28. AMOVA analysis based on ΦPT values showed 87% variation within and 13% among the population with statistical significance. Majority of the isolates from sugar beet showed nearby association within the population. There was significant number of cross crop clustering suggesting their broad pathogenicity. Isolates were grouped into three different clusters in UPGMA based cluster analysis using marker information. Interestingly, there was no specific geographical correlation between the isolates. PCA analysis showed randomized distribution among isolates from same geographical origin. Morphological characteristics showed crop-specific two distinct groups of isolates with few exceptions. While, genetic diversity showed two distinct group of isolates, one crop specific and one with wide pathogenicity. This information may help in molecular pathotyping of the pathogen for better disease management.

## 1.0 Introduction

*Rhizoctonia solani* is a polyphagous plant pathogen with worldwide distribution. It is a soilborne pathogen and known for severe plant diseases like collar rot, root rot, damping off and wire stem (Ogoshi 1996). The fungus survives on the infected plant debris and act as inoculation source for the susceptible crops like sugar beet (*Beta vulgaris* subsp. *vulgaris*) (Plyler-Harveson et al. 2011) dry beans (*Phaseolus vulgaris*) (Plyler-Harveson et al. 2011), potato (*Solanum tuberosum*) (Jung et al. 2012), and soybean (*Glycine max)* (Liu and Klein 2012). It is a major problem for the sugar beet and dry bean producers of western Nebraska. Total production acres of these crops are 45,500 and 140,000 – 200,00 acres respectively. *Rhizoctonia* root and crown rot in sugar beet and dry bean have reduced the yield significantly and has created problems in storage. It has been estimated that on average 2% of annual sugar beet yield loss is due to the *Rhizoctonia* rot, and even in some rare scenarios 30% - 60% to complete loss of the field had also been observed (Neher and Gallian 2011). In USA a minimum of In Nebraska, a total of 52% and 42% of yield reduction can be observed in case of dry bean variety viz. Great Northern Beans and Pinto beans respectively due to rhizoctonia rot (Harveson and Smith 2005).

*R. solani* occurs in varying degree of morphogenetic diversity. Cultural appearance, anastomosis, virulence, and physiology are different among different strains. Many scientific attempts have been made to categorize the *R. solani* isolates based on morphological, physiological and pathological differences. The most accepted grouping of *R. solani* is based on the formation of anastomosis or hyphal fusion (Carling 1996; Ogoshi 2003). There are now 14 anastomosis groups (AG), several of which are divided into subgroups (Carling et al. 2002). But presence of more than one AG and occasional loss of anastomosis ability always complicated identification and characterization of *Rhizoctonia sp.* Further, several studies have also indicated distinct pathogenesis even within the same AG groups (Ogoshi 2003; Dubey et al. 2012). Morphological characteristics are further influenced by the culture condition, which makes it difficult to characterize and categorize the isolates. The problem associated with characterization can be better addressed with the DNA-based molecular studies (TingDan et al. 2010; Dubey et al. 2012; Wang et al. 2013; Jaaffar et al. 2016b). Several DNA-based markers were used to analyze the genetic diversity of *R. solani viz.* genome sequence complementary analysis (Ceresini et al. 2002), Random Amplified Polymorphic DNA (RAPD) (Dubey et al. 2012; Wang et al. 2013; Shu et al. 2014), amplified fragment length polymorphism (AFLP) (Taheri et al. 2007), simple sequence repeats (Bernardes-de-Assis et al. 2009) and inter-simple sequence repeats (ISSR) (Sharma et al. 2005; Zheng et al. 2013; Wang et al. 2013; Zhou et al. 2014; Goswami et al. 2017).

ISSR marker was developed in 1994 and since then it is widely used for rapid differentiation among the closely related species. The technique involves the amplification of inter-region between two SSR region with a primer of 16 – 18 bp long and with a flanking region of nucleotides at 3’ or 5’ end. ISSR analysis is simple and less expensive than RAPD, and AFLP. It can be used to assess the genetic diversity of a large number of phytopathogen within relatively less time and with high reproducibility (Shu et al. 2014). Several researchers used dominant nature of the ISSR to establish the genetic variations and relationships among the *R. solani* isolates of different geographic province and within the same anastomosis group (Dubey et al. 2012; Wang et al. 2013; Shu et al. 2014).

Rhizoctonia root, stem, and crown rot is common in sugar beet and dry bean fields every year across western Nebraska. Our hypothesis was that isolates collected from two different crops could be different and isolates collected from the same crop across year and across geographic region could be different. Therefore, pathogen isolates were collected from both crops over years and were stored. The objective of this study was to determine the morpho-genetic diversity among *R. solani* isolates of sugar beets and dry beans from western Nebraska. The study was conducted over last ten-years of duration. The genetic diversity was assessed with 19 polymorphic ISSR markers.

## 2.0 Materials and Methods

### 2.1 Fungal isolates

Infected plants were collected from different regions of western Nebraska (**Fig. 1**). A total of 36 *R. solani* were isolated from the collected samples and used for this study. Twenty-seven were from sugar beet and nine isolates were from dry bean. Isolates were recovered from symptomatic diseased sugar beets and dry bean roots. For isolation, surface sterilized plant material was platted on potato dextrose agar (PDA) medium containing streptomycin antibiotic to reduce the opportunistic bacterial growth. Culture plates were incubated at 26 °C. Isolates were identified microscopically, and morphological features were recorded (**Table. 1**). Three control strains of *R. solani* of sugar beet origin were collected from Dr. Linda E. Hanson (USDA – ARS Sugar beet and Bean Research, East Lansing, Michigan) (**Table 1**). These strains were used for ISSR marker analysis to compare the genetic relatedness among the isolates and with known AG grouping. Population 1, 2, and 3, were used in this report to indicate the group of 27 sugar beet isolates, 9 dry bean isolates, and 3 control strains.

**Fig 1:**
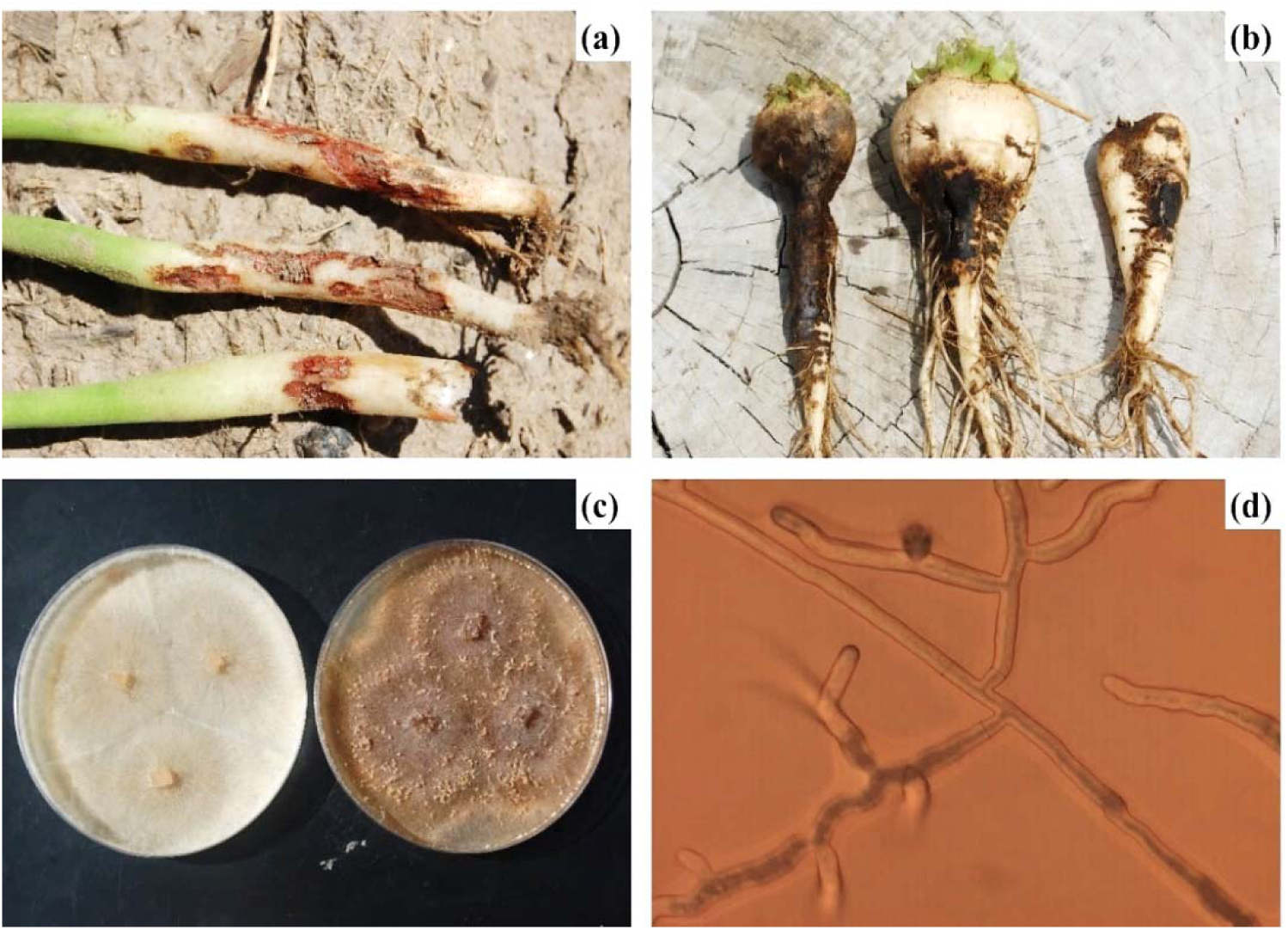
(a) Dry bean root rot (b) Sugar beet root rot (c) PDA culture plate (d) microscopic hyphal structure

**Table 1:**
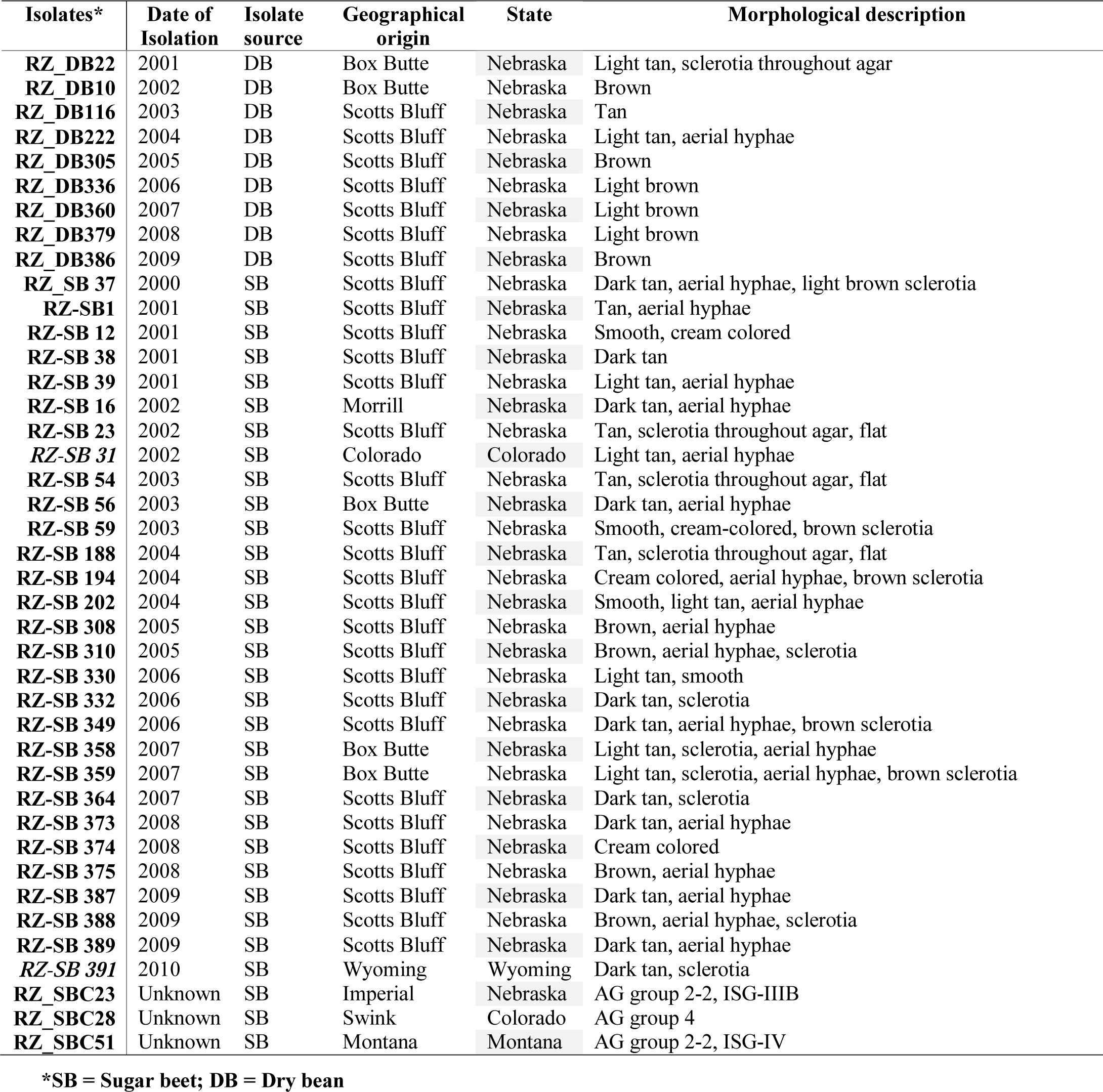
Details of 37 *Rhizoctonia solani* isolates with source, origin, year of isolation and morphological attributes.

| Isolates* | Date of Isolation | Isolate source | Geographical origin | State | Morphological description |
| --- | --- | --- | --- | --- | --- |
| **RZ_DB22** | 2001 | DB | Box Butte | Nebraska | Light tan, sclerotia throughout agar |
| **RZ_DB10** | 2002 | DB | Box Butte | Nebraska | Brown |
| **RZ_DB116** | 2003 | DB | Scotts Bluff | Nebraska | Tan |
| **RZ_DB222** | 2004 | DB | Scotts Bluff | Nebraska | Light tan, aerial hyphae |
| **RZ_DB305** | 2005 | DB | Scotts Bluff | Nebraska | Brown |
| **RZ_DB336** | 2006 | DB | Scotts Bluff | Nebraska | Light brown |
| **RZ_DB360** | 2007 | DB | Scotts Bluff | Nebraska | Light brown |
| **RZ_DB379** | 2008 | DB | Scotts Bluff | Nebraska | Light brown |
| **RZ_DB386** | 2009 | DB | Scotts Bluff | Nebraska | Brown |
| **RZ_SB 37** | 2000 | SB | Scotts Bluff | Nebraska | Dark tan, aerial hyphae, light brown sclerotia |
| **RZ-SB1** | 2001 | SB | Scotts Bluff | Nebraska | Tan, aerial hyphae |
| **RZ-SB 12** | 2001 | SB | Scotts Bluff | Nebraska | Smooth, cream colored |
| **RZ-SB 38** | 2001 | SB | Scotts Bluff | Nebraska | Dark tan |
| **RZ-SB 39** | 2001 | SB | Scotts Bluff | Nebraska | Light tan, aerial hyphae |
| **RZ-SB 16** | 2002 | SB | Morrill | Nebraska | Dark tan, aerial hyphae |
| **RZ-SB 23** | 2002 | SB | Scotts Bluff | Nebraska | Tan, sclerotia throughout agar, flat |
| **RZ-SB 31** | 2002 | SB | Colorado | Colorado | Light tan, aerial hyphae |
| **RZ-SB 54** | 2003 | SB | Scotts Bluff | Nebraska | Tan, sclerotia throughout agar, flat |
| **RZ-SB 56** | 2003 | SB | Box Butte | Nebraska | Dark tan, aerial hyphae |
| **RZ-SB 59** | 2003 | SB | Scotts Bluff | Nebraska | Smooth, cream-colored, brown sclerotia |
| **RZ-SB 188** | 2004 | SB | Scotts Bluff | Nebraska | Tan, sclerotia throughout agar, flat |
| **RZ-SB 194** | 2004 | SB | Scotts Bluff | Nebraska | Cream colored, aerial hyphae, brown sclerotia |
| **RZ-SB 202** | 2004 | SB | Scotts Bluff | Nebraska | Smooth, light tan, aerial hyphae |
| **RZ-SB 308** | 2005 | SB | Scotts Bluff | Nebraska | Brown, aerial hyphae |
| **RZ-SB 310** | 2005 | SB | Scotts Bluff | Nebraska | Brown, aerial hyphae, sclerotia |
| **RZ-SB 330** | 2006 | SB | Scotts Bluff | Nebraska | Light tan, smooth |
| **RZ-SB 332** | 2006 | SB | Scotts Bluff | Nebraska | Dark tan, sclerotia |
| **RZ-SB 349** | 2006 | SB | Scotts Bluff | Nebraska | Dark tan, aerial hyphae, brown sclerotia |
| **RZ-SB 358** | 2007 | SB | Box Butte | Nebraska | Light tan, sclerotia, aerial hyphae |
| **RZ-SB 359** | 2007 | SB | Box Butte | Nebraska | Light tan, sclerotia, aerial hyphae, brown sclerotia |
| **RZ-SB 364** | 2007 | SB | Scotts Bluff | Nebraska | Dark tan, sclerotia |
| **RZ-SB 373** | 2008 | SB | Scotts Bluff | Nebraska | Dark tan, aerial hyphae |
| **RZ-SB 374** | 2008 | SB | Scotts Bluff | Nebraska | Cream colored |
| **RZ-SB 375** | 2008 | SB | Scotts Bluff | Nebraska | Brown, aerial hyphae |
| **RZ-SB 387** | 2009 | SB | Scotts Bluff | Nebraska | Dark tan, aerial hyphae |
| **RZ-SB 388** | 2009 | SB | Scotts Bluff | Nebraska | Brown, aerial hyphae, sclerotia |
| **RZ-SB 389** | 2009 | SB | Scotts Bluff | Nebraska | Dark tan, aerial hyphae |
| **RZ-SB 391** | 2010 | SB | Wyoming | Wyoming | Dark tan, sclerotia |
| **RZ_SBC23** | Unknown | SB | Imperial | Nebraska | AG group 2-2, ISG-IIIB |
| **RZ_SBC28** | Unknown | SB | Swink | Colorado | AG group 4 |
| **RZ_SBC51** | Unknown | SB | Montana | Montana | AG group 2-2, ISG-IV |

\*SB = Sugar beet; DB = Dry bean

### 2.3 DNA Extraction

For DNA extraction fresh fungal culture was used. Five ¼ × ¼ inch plugs of agar were placed into 50 ml of sterile PDB and grown for 5 days at 26 °C. After the incubation period, mycelia were harvested by filtration through cheesecloth. The collected mycelia were lyophilized with liquid nitrogen and grounded into fine powder with mortar and pestle. The powder was transferred to 50 ml conical tubes containing 15 ml of CTAB extraction buffer (2% CTAB, 1.4 M NaCl, 20 mM EDTA, pH 8.0, 0.1 M Tris, pH 8.0, 0.4% B-mercaptoethanol) preheated to 65 °C. The samples were then incubated in a 65 °C water bath for one hour, with mixing at an interval of every 10 minutes. Samples were cooled for 10 minutes and 20 ml of chloroform/isoamyl alcohol (24:1) was added and mixed with each tube. The tubes were centrifuged at 3500 rpm for 20 minutes at 15 °C. The aqueous layer was transferred to a new tube and a double volume of chilled 95% ethanol was added to precipitate the DNA. The tubes were centrifuged at 3500 rpm for one minute and the supernatant was discarded. Pellets were washed with chilled 70% ethanol. After all ethanol was removed the dry DNA pellet was suspended in 1 ml of TE buffer (10 mM Tris-HCl and 1 mM EDTA, pH 8.0) and treated with 5 ul of RNAse (10mg/ml) at room temperature for one hour. DNA was quantified using the minigel method (Sambrrok et al. 1989) that compares band intensities with a standard lambda/HindIII DNA marker (Gibco BRL, Betheseda, MD) in a 0.8% agarose gel. DNA was diluted to a concentration of 25-50 ng/ul for use in polymerase chain reaction (PCR).

### 2.4 PCR amplification of ISSR

The 50 UBC primers screened in this study were obtained from eurofins genomics (Huntsville, Alabama). Nineteen (**Table 2**) were selected for analysis based on amplification profile (band intensity, quality and reproducibility in at least two independent replications) on two random isolates. PCR amplification was performed in 25 µl reaction mixture containing Promega 5× Green GoTaq Flexi Buffer, 2 mM MgCl2, 100 µm of dNTPs, 0.24 µM primer, 50-100 ng DNA, and 1 unit of *Taq* polymerase. The cyclic reaction was set at initial denaturation at 94 °C for 30 s, annealing at 50°C for 45 s, and an elongation at 72°C for 2 min (45 cycles). Final elongation was performed for 10 min. The PCR was completed on an Applied Biosystems Thermocycler 2720. PCR products were separated electrophoretically in 2% agarose gels including 0.5 µg/ml ethidium bromide and bands were visualized in the gel-doc system (Biorad, USA). ISSR gels were photographed using FOTO Analyst Express Electronic Imaging System (Fotodyne, Inc). Marker size range was determined by comparison with a 100bp DNA ladder (New England BioLabs). When scoring the gels, a marker locus was considered polymorphic if the band was not present in every isolate. Only clear DNA bands that were reproducible were scored. ISSR marker loci were designated by primer name.

**Table 2:**
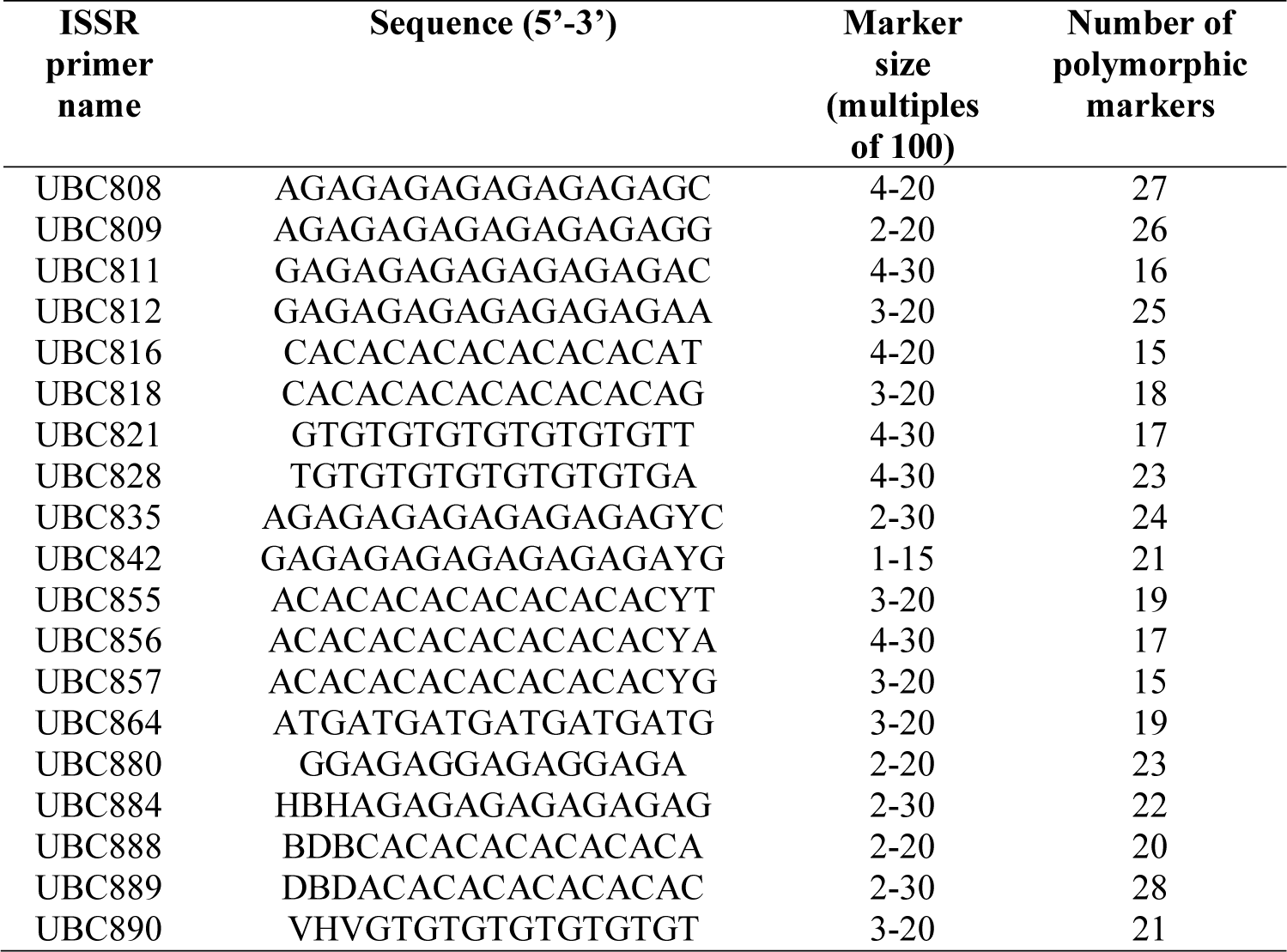
List of 19 ISSR markers used for the study

### 2.5 Data Analysis

A correlation matrix and dendrogram were prepared with unweighted pair group method with arithmetic mean (UPGMA) method to distinguish the isolates based on morphological characteristics. The categorical data of morphological traits were converted into 0, 1 matrix based on presence and absence. The dendrogram (method = UPGMA) and correlation plot (method = Pearson correlation) were prepared using R statistical software (package: pvclust and corrplot). Dominant polymorphic ISSR markers were scored based on presence and absence with 0 (absent) - 1 (present) matrix. Cluster analysis was carried out using UPGMA method and dendrogram was created by using R statistical software with a bootstrap value of 1000 with package pvclust (Suzuki and Shimodaira 2006). The package was used to compute two values: an approximately unbiased (AU) *p-*value based on multi step multi scale bootstrap resampling procedure (Shimodaira 2004) and a bootstrap probability (BP) *p-*value from ordinary bootstrap resampling (Efron et al. 1996). A significance threshold of α = 0.05 (95% confidence interval) was used in this approach. For a cluster with AU *p*-value > 0.95 (95%), the hypothesis that “the cluster does not exist” is rejected with significance level 0.05. The percentage of polymorphic loci (P), Shannon’s diversity index (*I*) and Nei’s gene diversity were calculated to estimate the genetic variation among the isolates using GenAlex 6.5 (Peakall and Smouse 2012). Genetic differentiation among populations was estimated by pairwise values of *Ф*_PT_. Analysis of molecular variance (AMOVA) was used to compute the genetic variation among and within the population. AMOVA calculations were performed in GenAlex 6.5. For AMOVA analysis, the control strains were not included, because of population size was three, which may create biases in the result. Principle component analysis was used to determined genetic differentiation among the isolates of different region using R statistical software (Package: FactoMiner, factoextra, and ggplot2). Morphological and genetic traits were correlated with R-stat to determine the diversity among and within the isolates of sugar beets and dry beans (Package: corrplot).

## 3.0 Results

### 3.1 Morphological Diversity

All the 38 isolates showed distinctive morphological variation in their appearance. Colony morphology varied from dark brown to light brown and light tan colored colonies **(Fig. 1),** sclerotia and presence of aerial hyphae (**Table 1**). Correlation analysis showed positive correlation within the isolates of same crops. The isolates from sugar beet showed positive correlation with sugar beet isolates and dry beans with dry bean isolates (**Fig. 2**). There was high positive correlation within the isolates of sugar beets, RZ_SB16, RZ_SB56, RZ_SB373, RZ_SB387, RZ_SB389, RZ_SB364, RZ_391, RZ_332 (r = 1.00 / 1.00, p <0.0001). Similarly, the isolates from dry beans R_DB10, RZ-DB305, RZ_DB386, RZ_DB116, RZ_DB336, RZ_DB360, RZ_DB379 showed significantly high correlation with each other (r = 1.00 / 1.00, p < 0.0001). Though most of the isolates showed morphological correlation within the isolates of same crop but there was also positive cross crop correlation with statistical significance but at lower degree. Isolates RZ_DB22 showed correlation with RZ_SB358 (r = 0.77/1.00, p < 0.01) and RZ_SB359 (r = 0.63/1.00, p < 0.05), RZ_DB305 with RZ-SB375 (r = 0.67/1.00, p < 0.05) (**Fig. 2**) (**Supplementary file: Table: T1**). There was no correlation between the isolates from same geographical origin and year of origin.

**Fig 2:**
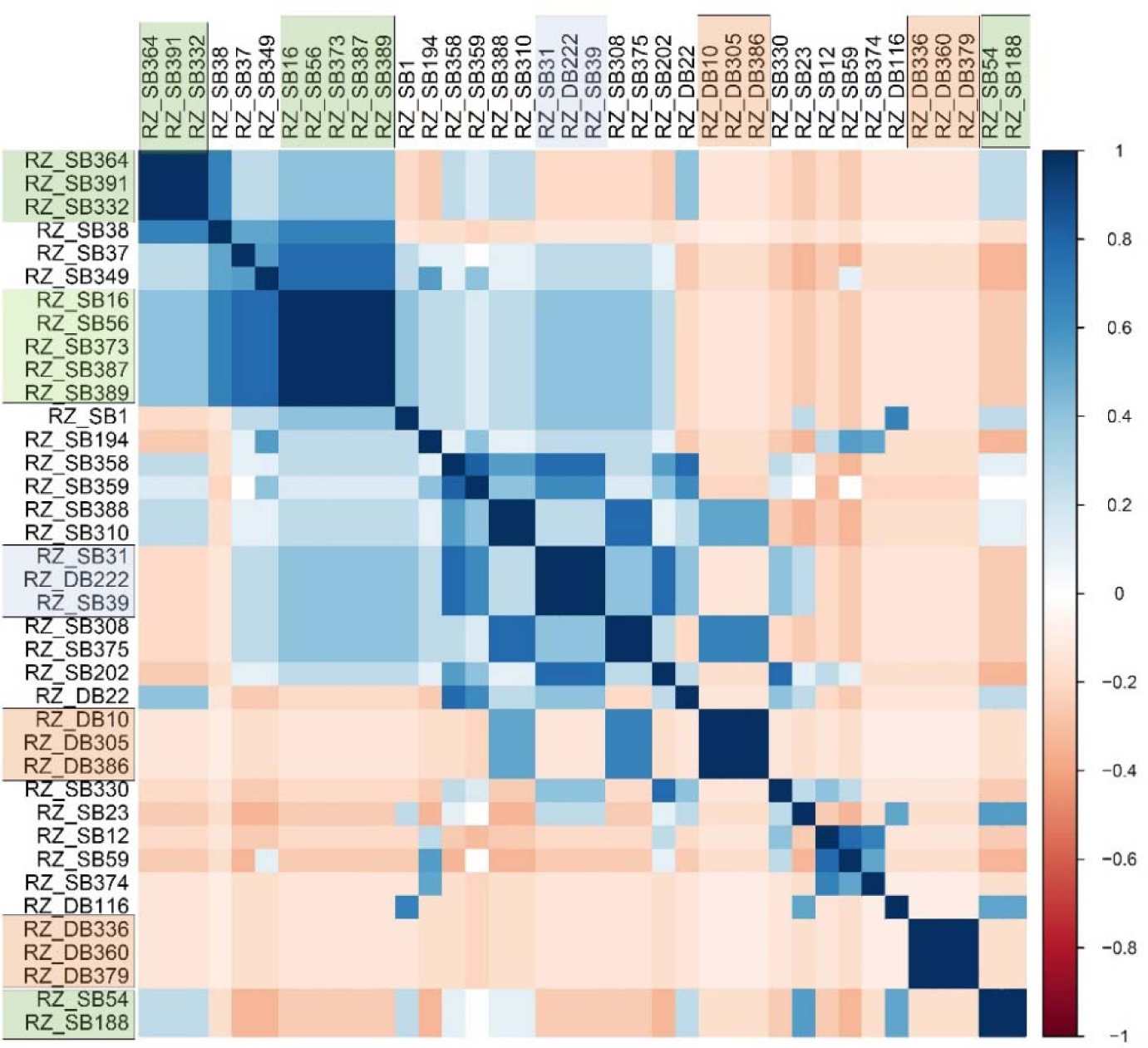
Correlation of the isolates based on their morphological characteristics. The isolates showed crop specific crop correlation. Sugar beet isolates were correlated with sugar beets and dry beans were correlated with dry beans with one exception. Correlation between sugar beets was highlighted with green boxed, orange box defines the correlation between dry beans and blue box was used for mixed correlation.

### 3.2 Genetic Diversity

A total of 50 UBC primers were screened and 19 primers were selected based on their 100% polymorphism index. A total of 396 alleles were identified from 41 isolates (representative gel image at Fig. 3). Average number of loci per primer was 20.84 and band size ranged from 1 – 3 kb. The primer UBC 889 produced the highest number of polymorphic loci (29) followed by UBC 808 (27), UBC 809 (26) and UBC 812(25) (**Table 2**). Shannon information index (*I)* varied from 0.235 – 0.280 with an average of 0.251. The percentage of polymorphic loci (% P) ranged 43.26% - 92.88%. The highest % polymorphic loci (92.88%) was observed within the dry bean isolates (population 2) (**Table 3**). Nei genetic distance within the population ranged from 0.033 – 0.083 with an average of 0.51.

**Fig 3:**
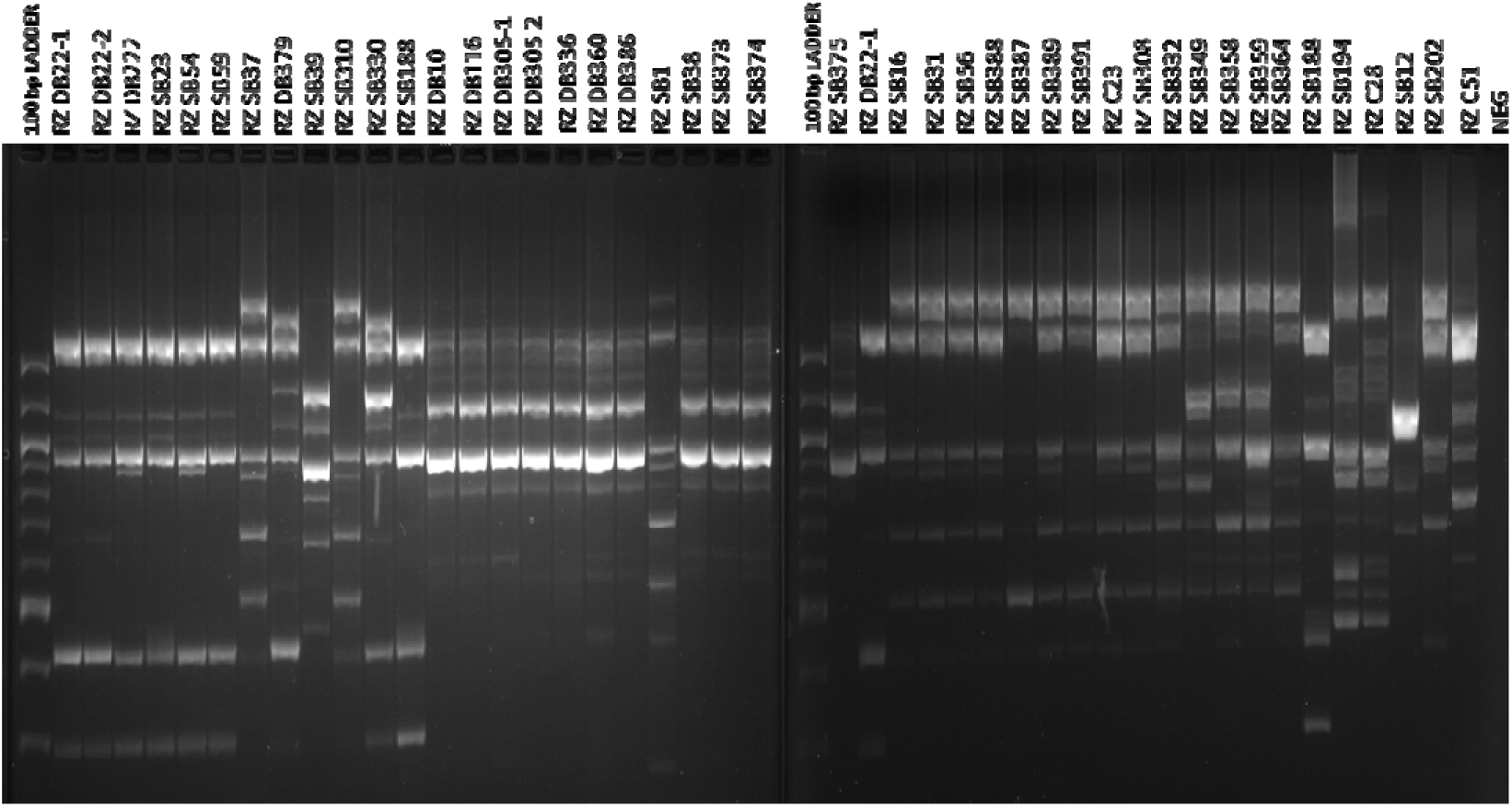
DNA marker profiles of *Rhizoctonia solani* isolates from sugar beets and dry beans with ISSR primer UBC809. Sugar beet and dry bean isolates are designated by RZ_SB and RZ_DB followed by a number.

**Table 3:** Allele diversity within the population

| Pop |  | N | Na | Ne | I | He | uHe | %P |
| --- | --- | --- | --- | --- | --- | --- | --- | --- |
| Pop1 | Mean | 8.000 | 0.980 | 1.269 | 0.240 | 0.159 | 0.169 | 48.35 |
|  | SE | 0.000 | 0.050 | 0.018 | 0.014 | 0.010 | 0.010 |  |
| Pop2 | Mean | 30.000 | 1.858 | 1.249 | 0.280 | 0.167 | 0.170 | 92.88 |
|  | SE | 0.000 | 0.026 | 0.014 | 0.010 | 0.008 | 0.008 |  |
| Pop3 | Mean | 3.000 | 0.891 | 1.259 | 0.235 | 0.156 | 0.187 | 43.26 |
|  | SE | 0.000 | 0.050 | 0.017 | 0.014 | 0.009 | 0.011 |  |

\*Na = No. of Different Alleles; Ne = No. of Effective Alleles = $1 / (p^2 + q^2)$ ; I = Shannon's Information Index = $-1 * (p * \ln(p) + q * \ln(q))$ ; He = Expected Heterozygosity = $2 * p * q$ ; uHe = Unbiased Expected Heterozygosity = $(2N / (2N-1)) * He$ [for Diploid Binary data and assuming Hardy-Weinberg Equilibrium, $q = (1 - \text{Band Freq.})^{0.5}$ and $p = 1 - q$ ]

Cluster analysis based on the UPGMA method had produced three distinct clusters. The first clusters majorly represented the isolates from the dry beans (RZ_DB 116, RZ_386, RZ_DB336, RZ_DB10, RZ_DB305, and RZ_DB360) showing their genetic similarity. However, four sugar beets isolates (RZ_SB373, RZ_374, RZ_375, and RZ_SB38) also showed significant similarities with the dry bean isolates in the first cluster. The second cluster included only sugar beet the isolates *viz.* RZ_SBC51, RZ_SB349, RZ_B358, RZ_SB37, RZ-SB1, RZ_SB389, RZ_SB391, RZ_SB16, RZ_SB56, RZ_SB338, RZ_SBC23, RZ_SB359, RZ_SB387, RZ_SB332, RZ_SB364. While the third cluster showed cross relation among the dry bean and sugar beet isolates (**Fig. 4**). Dry bean isolates like RZ_DB22, RZ_DB222, and RZ_DB379 showed genetic relatedness with RZ_SB330, RZ_SB188, and RZ_SB54 in the third cluster. Isolates like RZ_SB374 and RZ_SB375 (with au = 99% and bp = 97%) which were isolated in the year of 2008 from Scottsbluff showed similarity in genetic makeup. Isolates like RZ_SB332 (2005) showed similar genetics with isolates RZ_SB364 (2006) (au = 99% and bp = 96%) (#22).

**Fig 4:**
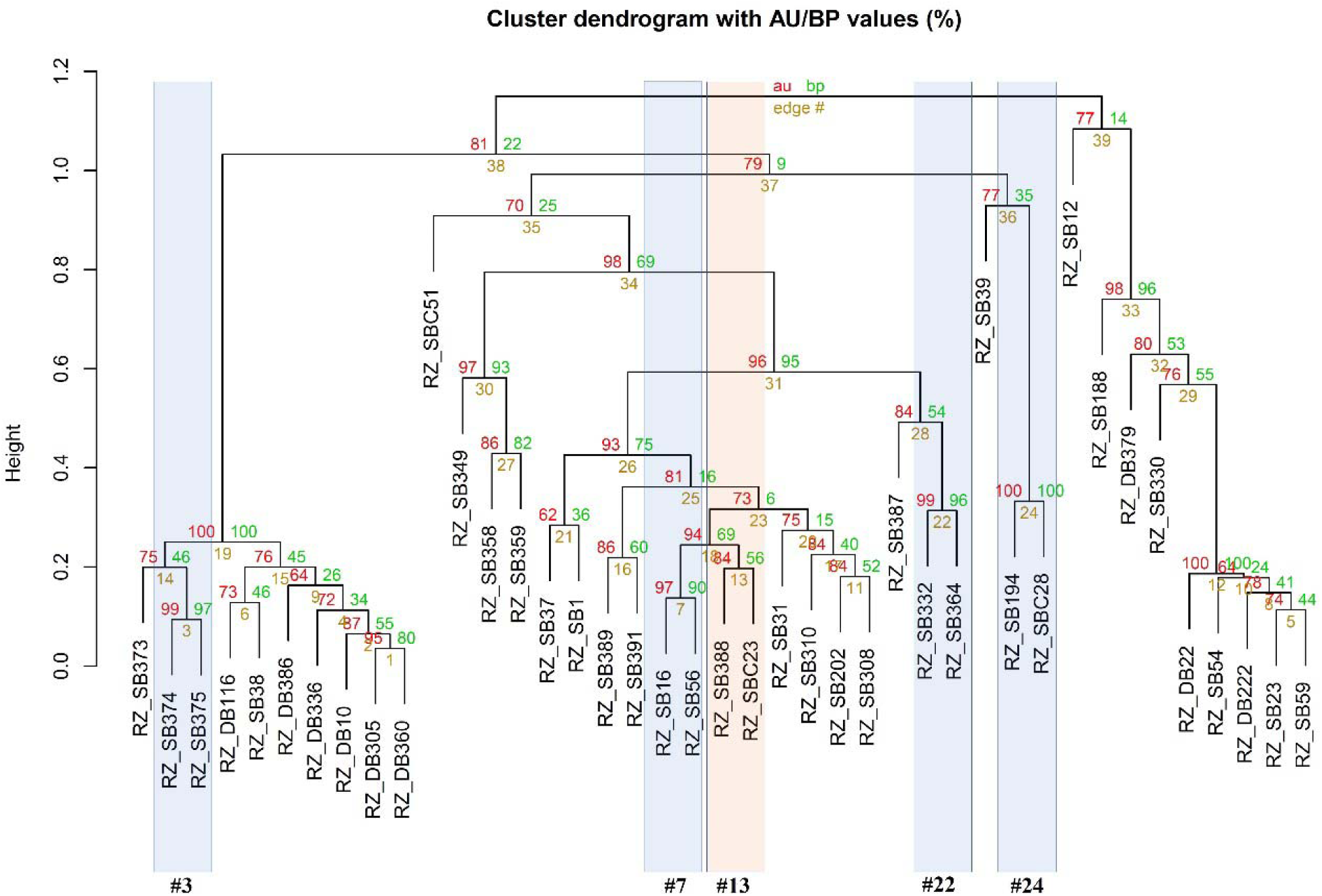
Dendrogram derived from the combined analysis of 19 ISSR prime for 41 Rhizoctonia solani with cluster method UPGMA. In the figure the values are represented as (%); where au = approximately unbiased p-value (red colored), bp = bootstrap probability (bootstrap value = 1000) (green colored) and #edge = number of the sub clusters (39 total clades) (yellow colored). Clusters with Au larger than 95% are strongly supported by the data. Clades (#clade number or edge#) were grouped based on the bootstrap identity and their characteristics. Blue highlighted clades are with high bootstrap identity (≥ 80%) and statistically significant (≥ 95%). Blue highlighted (#3, #7, #22, #24) clades are with high bootstrap identity (≥ 80%) and statistically significant (≥ 95%). Organge color highlighted clade is with high bootstrap value but statistically non-significant (#13).

The AMOVA analysis based on ΦPT values indicated that most of the genetic diversity occurred within the populations (87%, P <0.023) while variability among the population only contributed 13% (P <0.023) **(Table 4)**. Statistically significant genetic differentiation was observed among the isolates.

**Table 4:** Hierarchical distribution of genetic diversity among the population of *R. solani* from sugar beet and dry beans using AMOVA test

| Source | df | SS | MS | Estimated Variation | % of variation | ΦPT* | p |
| --- | --- | --- | --- | --- | --- | --- | --- |
| Among Pops | 2 | 219.93 | 109.97 | 7.29 | 13% | 0.14 | < 0.023 |
| Within Pops | 38 | 1787.53 | 47.04 | 47.04 | 87% |  |  |
| Total | 40 | 2007.46 |  | 54.33 | 100% |  |  |

\* ΦPT = $AP / (WP + AP) = AP / TOT$ (Key: AP = Est. Var. Among Pops, WP = Est. Var. Within Pops)

## Discussion

The genus *Rhizoctonia* is a diverse group of fungi which causes stem and root rot, foliar blights in many crops (Ogoshi 1996). *Rhizoctonia solani* Kuhn, a ubiquitous soil borne basidiomycetes, which causes diseases in many economically important crops like rice, potato, soybean, corn, sugar beet and dry beans (Sneh et al. 1991). In classical identification method, *R. solani* were characterized based on differences in pathogenicity, morphology and physiology (Sharma et al. 2005). In this study, a total of 36 *R. solani* was isolated from sugar beets and dry beans in a time span of 10 years from different location of western Nebraska, USA (**Table 1**). Studies on cultural characteristics revealed that the colony color of the different *R. solani* isolates varied from cream-colored to brown, dark tan to light tan in PDA culture plates with the production of areal hyphae and sclerotia with dark to light brown color (**Table 1**). The results showed a close agreement with other works (Takashi and Tadao 1978; Desvani et al. 2014; Jaaffar et al. 2016a).

Results of genetic analysis indicated a high degree of genetic diversity within the population (87%). Both the population of sugar beet and dry beans have unique genetic makeup which can be observed from the marker genotypes. The sugar beet and dry bean isolates are mostly conserved within each crop and formed distinct and different clusters in dendrogram analysis with few variations. Some of the sugar beet isolates also showed cross correlation with the isolates of dry bean (cluster 1 and cluster 3) (**Fig. 4**). This suggests relatedness among the population and wide pathogenicity spectrum of the group. AMOVA analysis also showed low variation among population (13%) compared to within the population (87%). PCA analysis and grouping based on marker genotypes also showed similar grouping patterns (Supplementary File: **Fig. S1**). It indicates however, the origin of *R. solani* for sugar beet and dry beans are same but there is certain degree of differentiation. The difference may have originated during the evolution and selection over pathogenicity. Similar results can be observed form the studies of Dubey et al., (2012), where the *R. solani* isolates were independent or did not corresponds to crops of origin (Stodart et al. 2007; Dubey et al. 2012).

Morphological classification of the isolates showed high degree differentiation among the isolates from sugar beet and dry beans. The morphological grouping mostly showed correlation within the isolates of same populations (**Fig. 2**). On differing, marker-based analysis showed cross correlations among the populations. Grouping of the isolates based on location, and isolation year over the marker data highlighted stimulating facts. The high bootstrap (97%) and p value (99%) between the isolates RZ_SB374 and RZ-375 indicates the same genetic origin. This matched with the isolate origin isolates since both were collected in same year (2008) and same place (Scottsbluff) (**Fig. 4**). It also indicates the chances of them being in the same pathovar group which couldn’t be morphologically differentiated. Isolates from consecutive years also showed high degree of genetic similarity defining the chances of same pathogens infecting the fields in the successive year. This also indicates their inoculum may have been present in the crop residues or soil from previous year, which are left untreated and produced the infection. Isolates like RZ_SB194 showed 100% bootstrap identity with control strain RZ_SBC28 of sugar beet (au =100%). This suggests genetic similarity and possibility of RZ_194 belonging to AG-Group 2-2 IIIB, which is a major anastomosis group responsible for sugar beet crown and root rot (Strausbaugh et al. 2011). Isolates RZ_SB388 and control strain RZ_SBC23 also showed genetic resemblance but with low bootstrap probability value (56%) and was not statistically significant (84%) (#13) (**Fig. 4**). It can be concluded, though there is some degree of similarity, but they are totally different pathovars.

The location based grouping of isolates on marker genotypes mostly showed independent nature of the isolates with random distribution. Scottsbluff with highest number of isolates showed correlation with all the isolates from other places and among themselves. Isolates from the scottsbluff showed high degree of similarity with the isolates of Box Bute, Morril, Imperial. Although sampling of this study was uneven with respect to crop and location, there was a mixed genetic population as noticed in cluster analysis, which distributed independent of their geographical locations (**Fig. 5**). Large group of isolates (Cluster 2) which showed genetic relatednes among and within the sugarbeet isolates only, while one group (Cluster 1 and Cluster 3) showed genetic relatedness among isolates from borth the crops. Among the 31 isolates from scottsbluff there is four distinct divisons. Among the four isolates of box bute two isolates showed correlation with their geographical location while other two were independent. Several studies reports this uneven relationship or no correlation between the place of origin and isolates of *R. solani* (Stodart et al. 2007; Zhou et al. 2009; Fiers et al. 2011; Dubey et al. 2012). Few studies do also reported geographical correlation (TingDan et al. 2010).

**Fig 5:**
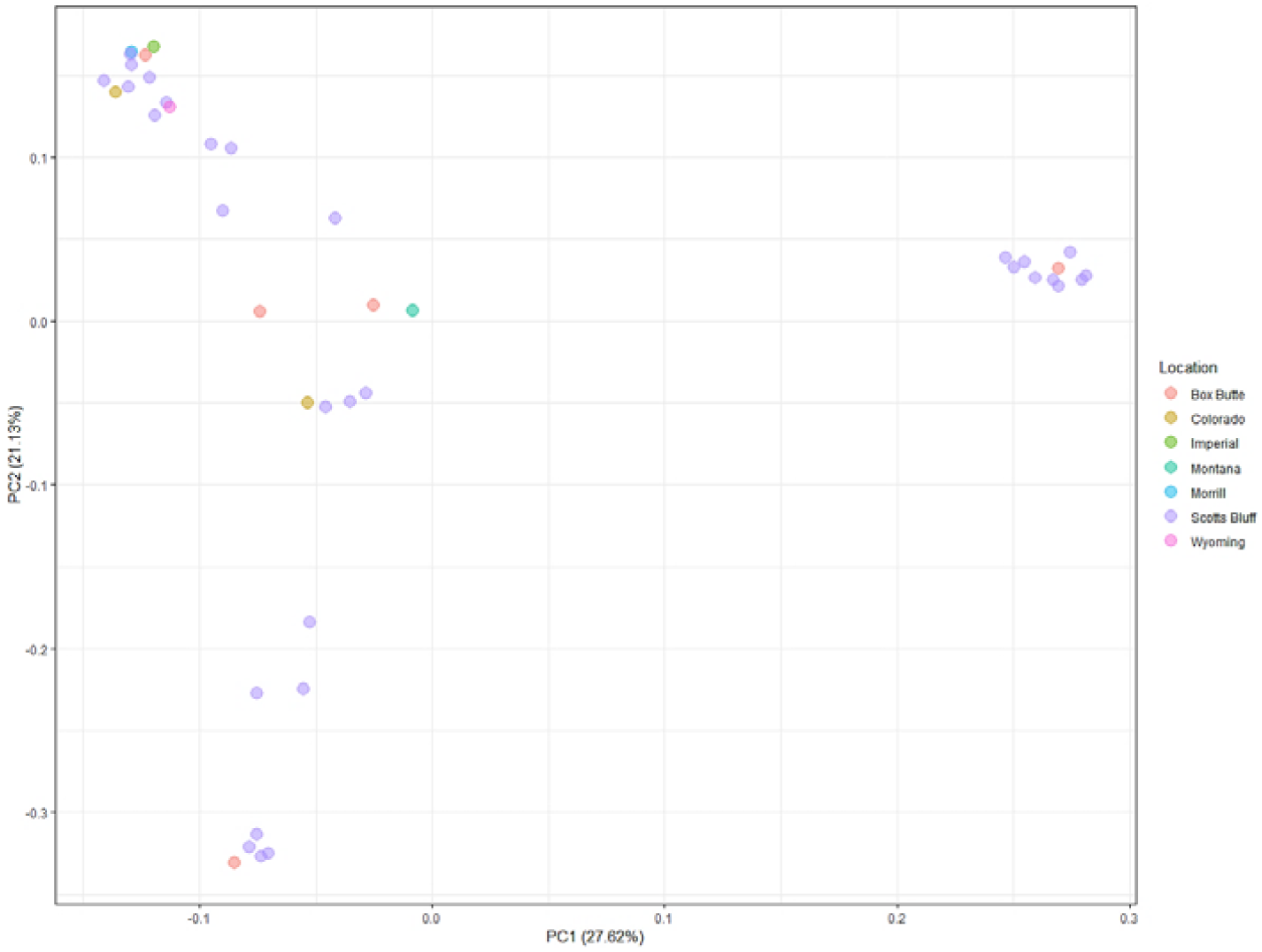
Location based distribution of the *R. solani* isolates based on marker data.

Relying only on morphological characteristics often may provide misidentification as the pathogens belonging to same group with similar mophological feature may have different pathogenocity. In this study, we observed morphological correlation showed distinct difference in between the isolates of sugar beet and dry beans, while genetic diversity showed significant differetiation. Therefore, taking morphological traits as sole identification method can results into biased grouping while the population are distinct or unique according to their genetic makeup. Thus, this study proved that marker information in conjucture with the morphological traits give better identifcation and characterization of intra and intergic gentic variations. For differentiation and characterizations of the *R. solani* from sugar beet and dry beans, ISSR marker was found suitable with the morphological features. This method can give a comprehesive estimation of genetic diversity of polyphyletic isolates where anastomosis or morphological features were not sufficient for conclusive characterization. Further studies based on ITS profiling and sequencing will be needed to provide more vivid knowledge on the genetic identity and variations with respect to crops and geographical origin. Identification and proper categorization of the pathogen will be helpful in desingning integrated disease management guidelines for sugar beet and dry beans of the mid western America.

## Acknowledgement

Authors would like to acknowledge Dr. Lisa E. Hanson, USDA Research Scientist, for providing the control strains for *R. solani.*

## Authors Contribution

DS & RH conceived the idea and designed the study. SD, & TB performed the experiments, data analysis and prepared the manuscript. DS & RH reviewed and prepared the final draft. All authors have read and approved the final manuscript.

